# Separation and Loss of Centrioles From Primordidal Germ Cells to Mature Oocytes in the Mouse

**DOI:** 10.1101/339572

**Authors:** Calvin Simerly, Marion Manil-Ségalen, Carlos Castro, Carrie Hartnett, Dong Kong, Marie-Hélène Verlhac, Jadranka Loncarek, Gerald Schatten

## Abstract

Oocytes, including those from mammals, lack centrioles, but neither the mechanism by which mature eggs lose their centrioles nor the exact stage at which centrioles are destroyed during oogenesis is known. To answer questions raised by centriole disappearance during oogenesis, using a transgenic mouse expressing GFP-centrin-2 (GFP CETN2), we traced their presence from e11.5 primordial germ cells (PGCs) through oogenesis and their ultimate dissolution in mature oocytes. We show tightly coupled CETN2 doublets in PGCs, oogonia, and pre-pubertal oocytes. Beginning with follicular recruitment of incompetent germinal vesicle (GV) oocytes, through full oocyte maturation, the CETN2 doublets separate within the pericentriolar material (PCM); concomitantly, a rise in single CETN2 pairs is identified. CETN2 dissolution accelerates following meiosis resumption. Remarkably, a single CETN2 pair is retained in the PCM of most meiotic metaphase-I and -II spindle poles. Partial dissolution of the CETN2 foci occurs even as other centriole markers, like Cep135, a protein necessary for centriole duplication, are maintained at the PCM. Furthermore, live imaging demonstrates that the link between the two centrioles breaks as meiosis resumes and that centriole association with the PCM is progressively lost. Microtubule inhibition shows that centriole dissolution is uncoupled from microtubule dynamics. Thus, centriole doublets, present in early G2-arrested meiotic prophase oocytes, begin partial reduction during follicular recruitment and meiotic resumption, much later than previously thought.

## Introduction

Centrioles, found at the poles of mitotic spindles, are vital for reproduction and development. Long thought to be contributed by the sperm during fertilization and lost during fetal oogenesis, they are essential in innumerable processes (rev, Schatten, 1994). Indeed, centriole defects appear as the root causes of a broad set of diseases, ranging from blindness and cancers through microcephaly and ciliopathies (Boveri, 1914; Bettencourt-Dias et al, 2011). Centrioles are often surrounded by the pericentriolar material (PCM), and together, the two structures define the canonical centrosome, the cell’s major microtubule organizing center (MTOC; Bettencourt-Dias et al, 2011).

In most mammals, haploid female gametes produced during oogenesis lose their centrosomes, although the mechanism of when and how remains elusive (Manandhar et al, 2005; Loncarek and Khodjakov, 2009). Most studies on centrosome reduction in gametes involve ultrastructural observations (Szollosi et al, 1972; Santhanthan et al, 2006; Manandhar et al, 2005). In humans, centrioles have been detected in fetal oogonia at 13-15 weeks post-gestation and within early growing oocytes (Sathananthan et al, 2000). However, centrioles have not been found in fully grown germinal vesicle (GV)-stage oocytes, and the metaphase-I and -II spindles formed after meiotic resumption are anastral, barrel-shaped structures with spindle poles devoid of centrioles or PCM (Sathananthan et al, 2006). In mice, ultrastructural and marker tracing have identified intact centriole pairs in fetal oogonia and early post-natal stage (P4) mouse primordial oocytes (Kloc et al, 2008; Lei and Spradling, 2016; rev Pepling, 2016). In later, preovulatory stages, growing mouse oocytes apparently lose centrioles (Calarco, 2000) while maintaining dispersed acentriolar PCM throughout the cytoplasm.

As the oocyte reaches maturity and competency to enter meiosis, a perinuclear MTOC, composed of PCM constituents such as γ-tubulin and pericentrin, gradually enlarges near the GV nucleus (Carabatsos et al, 2000; Dumont et al, 2007; Łuksza et al, 2013). Upon meiotic resumption, the acentriolar PCM fragments along the GV nucleus, mediated by PLK1, which releases the centriole adhesion protein cNAP1 (centrosomal Nek2-associated protein-1; Mayor et al, 2000) and then is stretched and fragmented by BicD2-anchored dynein in a microtubule-dependent manner (Łuksza et al, 2013). Finally, KIF11 mediates further MTOC fragmentation to allow segregation of PCM material to opposing spindle poles (Clift and Schuh, 2015). The kinases Aurora A and PLK4 also enhance microtubule growth and first meiotic spindle assembly as chromosomal divisions ensue (Bury et al, 2017). The arrested mouse metaphase-II spindle is anastral and acentriolar but maintains assembled PCM material at the spindle poles and within distinct cytoplasmic foci (Maro et al, 1985; Schatten et al, 1985, 1986; rev, Schatten, 1994). Interestingly, the mouse sperm does not contribute a centriole at fertilization (Woolley and Fawcett, 1973; Manandhar et al, 1999; Simerly et al, 2016), and zygotes rely on convergent cytoplasmic PCM and kinesin-5 to progress through mitotic divisions during early development until the blastocyst stage, when centrioles reappear at the spindle poles (Guet-Hallonet et al, 1993; Palacios et al, 1993; Howe and FitzHarris, 2013a, b).

The most prominent permanent core components found, nearly universally, in the centriole and within the centrosomes are centrin, pericentrin, and γ-tubulin. Centrin is an EF-hand calcium-binding protein found in the lumen of assembled centrioles (Baron et al, 1992). Centrins are required for basal body formation and positioning of the spindle pole body in yeast, algae, and ciliates (Salisbury et al, 2002; Salisbury, 2007). Mammals express four centrin genes (CETN1-4), but their cellular functions are not known (Friedberg, 2006; Bornens and Azimzadeh, 2007). γ-tubulin is the tubulin isoform responsible for serving as the MTOC (Oakley et al, 2015) and is a component of the γ-tubulin ring complex (γ-TuRC; Kollman et al, 2015). Pericentrin is a conserved coiled-coil PCM scaffolding protein that complexes with γ-tubulin and other proteins to initiate microtubule nucleating activity and cell cycle regulation (Doxsey et al, 1994; Delaval and Doxsey, 2010).

Centrioles have been reliably traced dynamically with transgenic reporter green fluorescent protein (GFP)-labeled centrin, including GFP-centrin-2 (GFP CETN2; White et al, 2000; Piel et al, 2000; D’Assoro et al, 2001; Kuriyama et al, 2007; Zhong et al, 2007; Balestra et al, 2013). A stable transgenic mouse strain that constitutively expresses GFP CETN2 in every cell of the body, including gametes, has been generated and shown to be a reliable probe for tracing centriole behavior in a variety of organs in the mouse (Higginbotham et al, 2004; Simerly et al, 2016). These transgenic mice, expressing GFP CETN2, appear nearly normal physiologically when compared to wild-type mice (Higginbotham et al, 2004; Howe and FitzHarris, 2013b).

Here, using this transgenic mouse constitutively expressing GFP CETN2 to trace centrioles and γ-tubulin to track centrosomes, along with microtubule and DNA probes, we found centriole pairs in somatic cells (e.g., cumulus and stromal cells) as well as in oogonia from fetal ovaries persisting in immature oocytes in the adult ovaries. As maturing oocytes transition into full competency to resume meiotic maturation and chromosome-reductional divisions, these pairs begin to disassemble, although they maintain the ability to nucleate microtubules throughout meiosis. Surprisingly, GFP-centrin-tagged structures are visible at meiotic spindle pole PCM at metaphase-I and -II spindles. Yet, following both GFP-centrin and PCM in living oocytes shows that their association is weak and that, in oocytes, the CETN2 sites do not retain the capacity to organize the PCM as meiosis I progresses as they do in mitotic cells. These foci comprise the largest, best organized PCM in the cytoplasm, and drug recovery from nocodazole microtubule disassembly shows prominent microtubule nucleation from these GFP CETN2-containing PCM foci after drug reversal. Collectively, this study suggests that centriole remnants persist in fully grown oocytes and that the dogma regarding the destruction of the maternal centrioles in early oogenesis may need to be reconsidered.

## Results

### PRIMORDIAL GERM CELLS AND OOGONIA IN FETAL OVARIES CONTAIN GFP-CENTRIN2-LABELED CENTRIOLES

Since centriole elimination is reported to occur in fetal stages (Manandhar et al, 2005), we first investigated primordial germ cells (PGCs) in the nascent gonads in GFP CETN2-expressing CB6F1 females, with sex identified by y chromosome PCR (see Methods and Materials), and traced their history through fetal oogonia formation in developing ovaries prior to birth. Isolated PGCs from e11.5 postcoitus gonadal tissues were identified from undifferentiated or somatic cells by the expression of the germ line cell surface marker Stage-Specific Embryonic Antigen-1 (SSEA-1; Fig. 1; Fenderson et al, 2006). Initial findings on this transgenic mouse reported GFP CETN2 expression from e14.5 forward (Jackson Laboratory Mouse Database Description at https://www.jax.org/jax-mice-and-services#; Higginbotham et al, 2004), so we were initially surprised to find that all e11.5 PGCs also contained GFP CETN2-labeled centrioles within microtubule arrays, whether mitotic or interphase (Fig. 1A-D). When PGCs stopped mitotic divisions, and entered pre-meiotic G2 arrest around e12.5 post-coitus, twin pairs of GFP CETN2 foci, in association with microtubules, were identified in SSEA-1-positive e14.5 and e18.5 oogonia, as well as in surrounding somatic cells (Fig. 1E-H). We conclude that GFP CETN2 centrioles are present and functional throughout fetal development, including their presence in female primordial germ cells.

**Figure 1.**
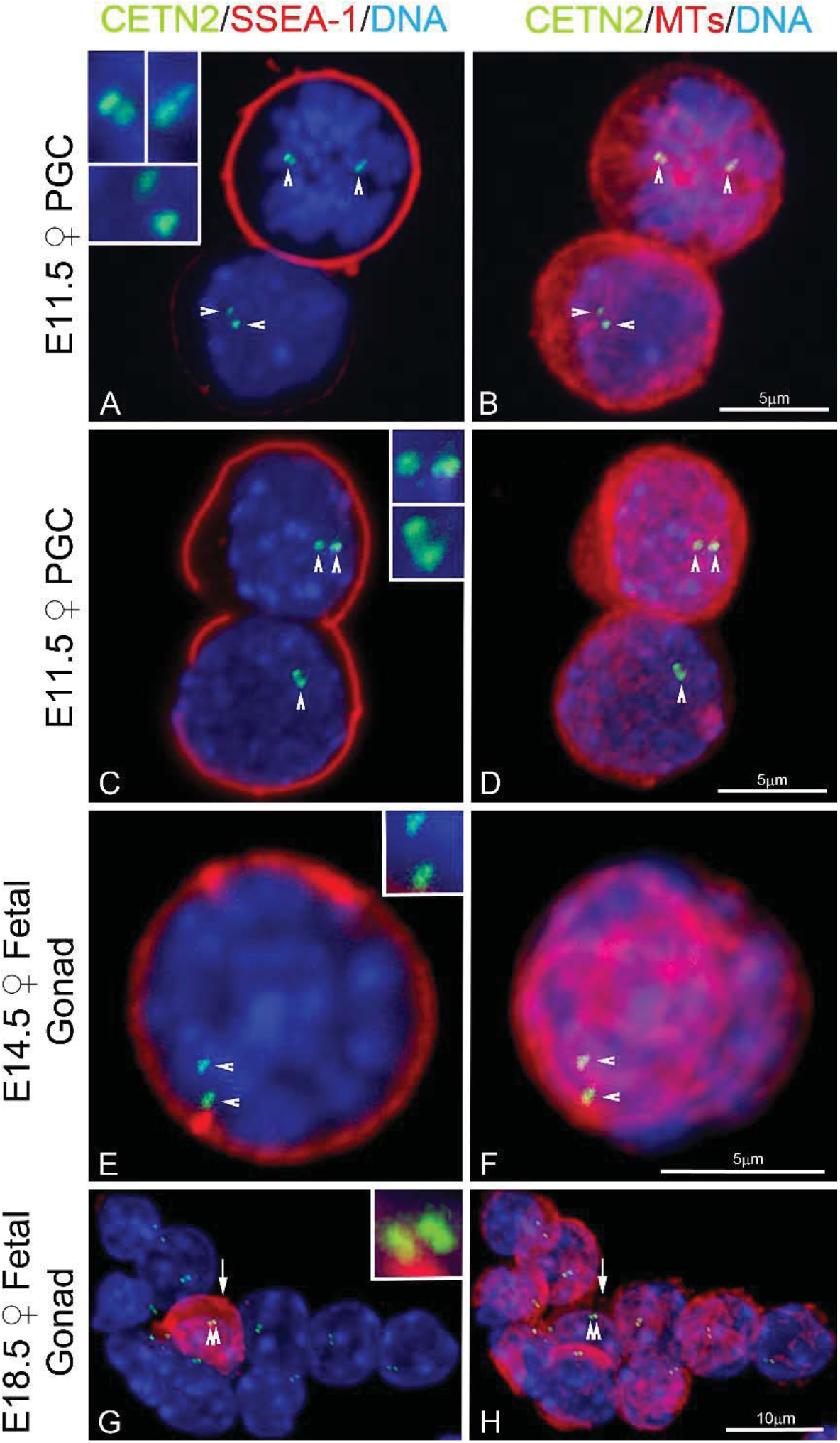
GFP CETN2 centriole detection in isolated mouse primordial germ cells and fetal oogonia. (**A-B**): isolated primordial germ cells (PGCs) from an e11.5 dpc female gonad. A: upper mitotic cell (blue, DNA) shows the presence of the germ cell marker SSEA-1 (red) at the cell periphery, along with two pairs of GFP CETN2 foci (inset) on opposite sides of the condensed prometaphase chromosomes (green, arrowheads). The lower somatic interphase gonadal cell lacks SSEA-1, with a single pair of GFP CETN2 focus (inset) adjacent to the nucleus (green, arrowheads). All GFP CETN2 centrioles are associated with microtubules, either at the center of microtubule asters at each nascent spindle pole (B: upper cell, red) or assembled on the nuclear surface connected to the more robust cortical microtubule interphase network (B: lower cell, red). (**C-D**): a divided e11.5 dpc primordial germ cell with SSEA-1 at the cell periphery of both daughter cells (C: red) and expressing GFP CETN2 centrioles (C: green, arrowheads) adjacent to the nuclei (C: blue, DNA). Microtubule assembly is observed only from the top daughter cell (D: red). (**E-F**): early mouse oogonium from an e14.5 dpc gonad with cortical SSEA-1 (E: red) and GFP CETN2 centriole pairs (E: green, arrowheads; blue, DNA). Cortical microtubule bundles are pronounced at the site of the GFP CETN2 centriole pair (F: red, microtubules, arrowheads). (**G-H**) Cells isolated from the sex cord of an e18.5 dpc fetal gonad, showing a single oogonial cell labeled with SSEA-1 (G: red, arrow) with a pair of GFP CETN2 centrioles (G: green, double arrowheads) at the nucleus (blue, DNA) but without apparent microtubule assembly (H: red, double arrowheads). The remaining somatic interphase gonadal cells are SSEA-1-negative but express GFP CETN2 centriole pairs (G: green foci) at their nuclear surfaces near the site of interphase microtubule assembly (H: red, microtubules). Confocal images of GFP CETN2 (green) are counterstained with antibodies to SSEA-1 (A, C, E, G: red), microtubules (B, D, F, H: color assigned red) and DNA (blue). Insets in A, C, E, G: details of GFP CETN2 centrioles. dpc: days post-coitus. Bars as marked.

### DIMINISHED GFP CETN2 PUNCTATE FOCI, PERHAPS CENTRIOLE REMNANTS, ARE TYPICALLY OBSERVED IN METAPHASE-I AND -II SPINDLE POLES

We traced the fate of the GFP CETN2-labeled structures during meiotic maturation. At meiotic resumption, separated GFP CETN2-labeled pairs embedded in γ-tubulin ribbons were observed at the assembling spindle poles (Fig. 2A-G). Cumulus cells, serving as an internal somatic cell control, also showed GFP CETN2-labeled centriole pairs within γ-tubulin PCM, including at the spindle poles of a rare bipolar mitotic cell (Fig. 2A, 2C: arrowheads). Using correlative light and electron microscopy (CLEM), we investigated whether classical 9-triplet microtubule organization could be identified in oocytes at this stage (Suppl Fig. S1). Analysis revealed linear MTOC-like structures with associated GFP CETN2 foci (Supple Fig. S1B, inset) in live oocytes. EM sectioning through the imaged site, however, showed only linear MTOC structures and multi-vesicular PCM aggregates (Calarco, 2000; Supple Fig. S1B-F), but no classical centrioles with microtubule walls, despite careful examination of hundreds of 80-nm-thick sequential serial sections. Typical canonical centrioles were identified in adhering cumulus cells attached to the same oocyte (Fig. 2E, inset; Suppl Fig. S1G-I). Collectively, the data suggest that significant oocyte centriole dissolution begins with meiosis onset. Altered oocyte centrioles still display GFP CETN2 labeling but are either too fragile to survive EM processing protocols or have undergone partial disassembly. This observation is supported by finding classical centrioles with 9+0 triplet microtubules in the surrounding somatic cumulus cells.

**Figure 2.**
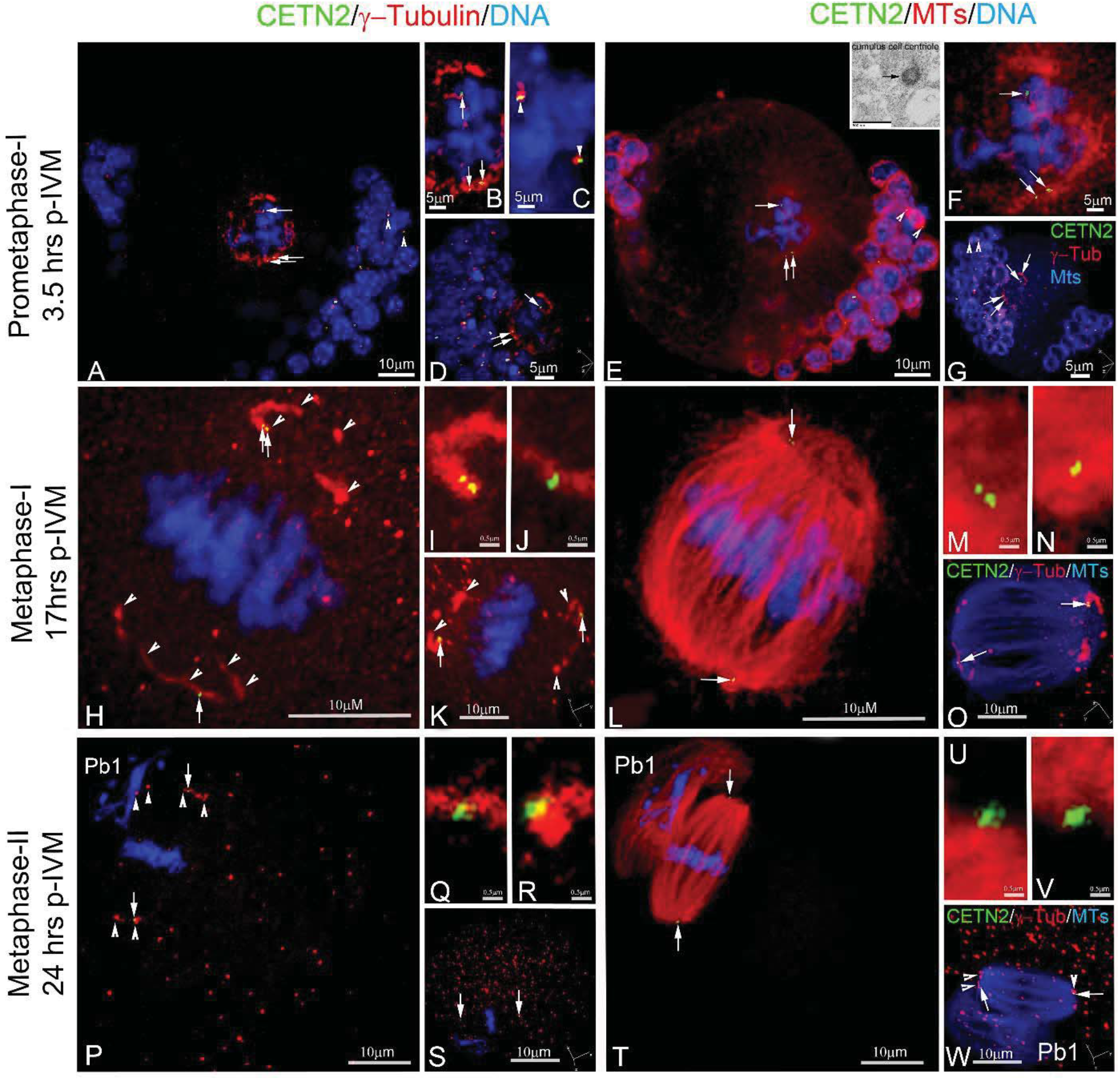
GFP CETN2 foci associate with developing spindle poles during meiotic maturation up to and including metaphase-II arrest. (**A-G**): selected z-projection of a prometaphase-I oocyte during early spindle assembly. GFP CETN2 foci are visible at opposite developing spindle poles (A: green, arrows), embedded in an expanded ribbon of y-tubulin (A: red; B: magnified view) surrounding the condensed bivalents (blue). Cumulus cells at the cell periphery (blue) also express GFP CETN2 centrioles (green) surrounded by y-tubulin (red), including a cell at mitotic metaphase (A: arrowheads; C: magnified view of cumulus metaphase spindle with GFP CETN2 centrosomes). Spindle microtubules (E: red; F: magnified view) assemble from the GFP CETN2: y-tubulin centrosomes. Inset, E: EM cross-section of a cumulus cell centriole with the canonical 9 + 0 triplet microtubule pattern (arrow; see also Suppl Fig. S1G-I). D, G: rotational views around the developing prometaphase-I spindle (axis, lower right in panels). (**H-W**): z-projections in metaphase-I-(H-O) and metaphase-II-arrested (P-W) meiotic spindles. GFP CETN2 foci (H, P: green, arrows) are identified at opposite poles of the bipolar spindles (L, T: red, microtubules), with aligned chromosomes (blue) and y-tubulin ribbons that encircle each pole (H, P: red, arrowheads). Pb1: the first polar body, showing no GFP CETN2 foci (green) despite y-tubulin (closed arrowheads) and disarrayed microtubule assembly (T, red). I, J, M, N: magnified views of GFP CETN2 foci (green) at metaphase-I spindle poles within y-tubulin ribbons (I, J: red) or assembled microtubules (M, N: red). K, O: metaphase-I spindle rotational views (axis orientations, lower right in panels). Q, R, U, V: magnified views of the metaphase-II spindle GFP CETN2 foci (green) embedded in y-tubulin (Q, R: red) or in relation to assembled spindle microtubules (U, V: red). S, W: metaphase-II spindle rotations (axis, lower right in panels). The extraneous red foci in panels K, S, and W appear to be non-specific binding of the polyclonal AK15 y-tubulin antibody. All images are of directly expressing GFP CETN2 oocytes (green), counterstained for y-tubulin (red), microtubules (cy5, color assigned red), and DNA (blue). Bars in μm.

Support for oocyte centriole alterations occurring during meiotic resumption was also derived from measuring changes in PCM and GFP CETN2 foci area sizes (in μm^2^) during maturation (Suppl Fig. S2E). After germinal vesicle breakdown (GVBD) and through complete maturation to metaphase-II arrest, we measured increased γ-tubulin area as the GV-associated MTOC foci were stretched into expanded ribbons and fragmented, as previously reported (Łuksza et al, 2013; Clift and Schuh, 2015). However, significantly decreased GFP CETN2 areas were measured after GVBD, suggesting structural changes in oocyte centrioles with diminishing GFP CETN2 detection. Interestingly, areas for GFP CETN2 centrioles in somatic cumulus cells were consistent in primordial follicles, growing antral follicles, and surrounding mature GVs after follicular release (0.47 ± 0.17, 0.42 ± 0.23, and 0.43 ± 0.19 μm^2^, respectively). Despite reduced areas in GFP CETN2 foci after meiotic resumption, most spindle poles in metaphase-I and -II oocytes still contained GFP CETN2 foci at the ends of γ-tubulin short segments or foci on the outer edges of the barrel-shaped spindles (Fig. 2H-W, Suppl Fig. S2F).

Tracking GFP CETN2 foci in first and second meiotic spindles showed many localization patterns. Most meiotic spindles contained single GFP CETN2 foci at the extreme edges of the barrel-shaped spindle (Fig. 3 and Suppl Fig. S3A-G), unlike centrosomes in somatic cell spindles that reside more centrally in the fusiform spindle pole. We also found GFP CETN2 foci within the spindle lattice near the aligned bivalents, either with or without γ–tubulin PCM (Suppl Fig. S3H-V). In metaphase-II-arrested oocytes, a minority of GFP CETN2 foci were observed in the extruded first polar body (Pb1) or in the microtubule-based asters (cytasters; Maro et al. JCB 1985; Schatten et al, 1985,1986) in the cytoplasm (Suppl Fig. S4). However, the majority of GFP CETN2 foci were found in association with meiotic spindles throughout maturation to metaphase-II arrest (Suppl Fig. S2F). Taken together, our analyses suggest that centrioles show significant reduction during meiotic progression and could not be identified as traditional centrioles with 9-fold symmetry in EM serial sections, consistent with centriole dissolution prior to the onset of meiosis. However, GFP CETN2 foci identified in fetal stages survive in mature oocytes and are found in γ-tubulin-containing meiotic spindle poles through metaphase-II arrest following meiotic resumption. GFP CETN2 foci are found in atypical spindle pole positions relative to traditional localization of centrioles in somatic cells.

**Figure 3.**
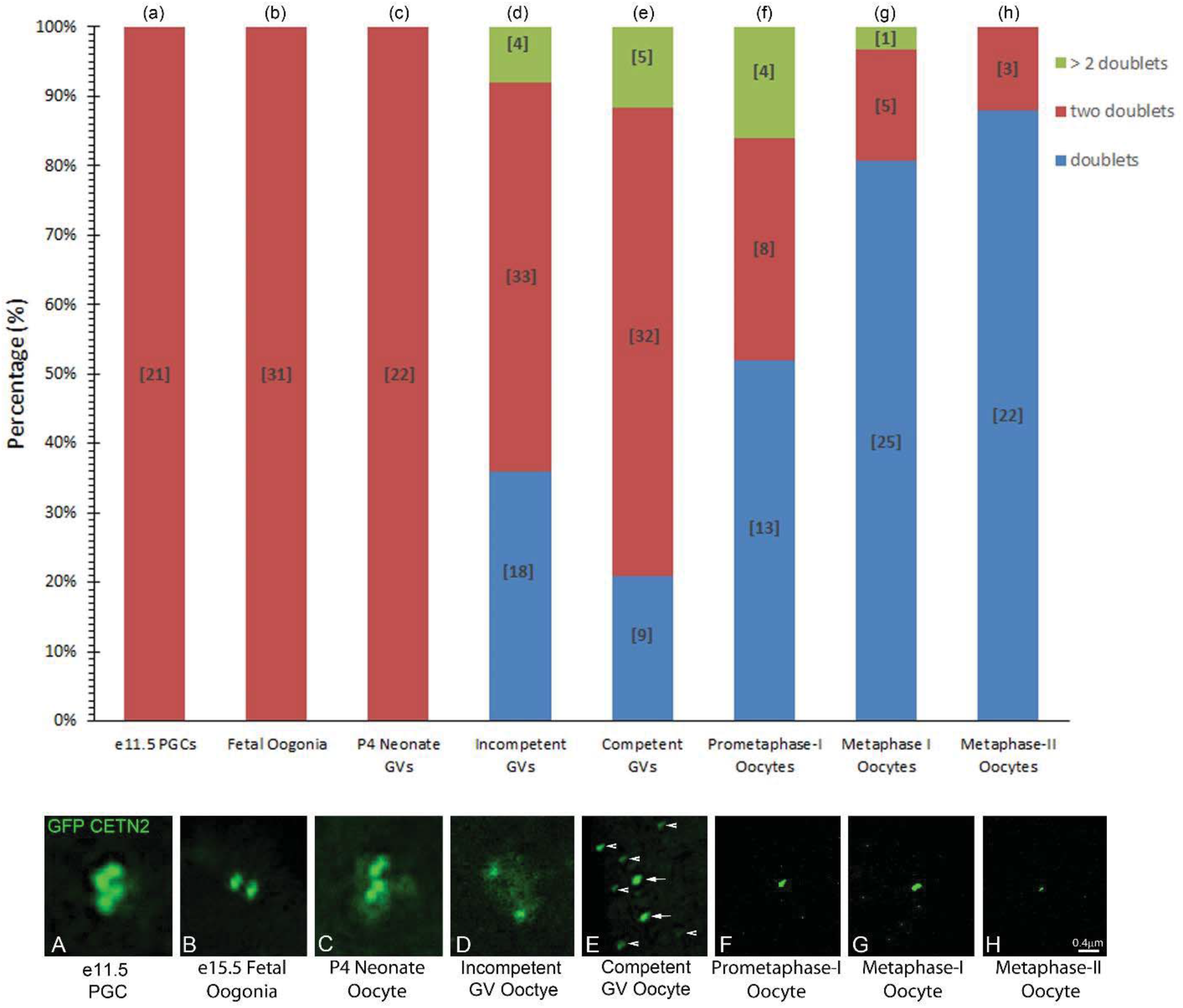
Tracking GFP CETN2 centriole loss from PGCs through maturation to metaphase-II arrest. Nested bar depiction of oocyte populations in PGCs, oogonia, adolescent (P4) primary oocytes, various adult oocytes during follicular growth, and meiotic oocytes showing the percentage of GFP CETN2 foci as doublets (1 pair), two doublets (2 pair), or > two doublets (i.e. more than 3 doublets) at each stage. Tightly apposed GFP CETN2 doublets in association with PCM are observed in all e11.5 PGCs, fetal stage oogonia (e14.5 and e18.5), and neonate early oocytes prior to sexual maturity (graph: a-c, red bars; image panel: 3A-C, green). In adult ovaries, however, follicular recruitment under the influence of sex hormones alters the pattern of GFP CETN2 doublets during oocyte growth to full maturity, reducing both the number of doublets (graph: d-e, red bars) and their tight association (image panel: D-E, green; arrows point to GFP CETN2 doublets; arrowheads: GFP aggregates). GFP CETN2 doublets continue to decline with the onset of meiotic maturation (graph: f-h, red bars), with a significant increase in GFP CETN2 doublets or single pairs within the PCM at a single spindle pole in metaphase-I and -II oocytes (graph: f-h, blue bars; image panel: F-H, green). Other GFP CETN2 configurations include multiple doublets observed in immature, fully grown, and early prometaphase-I oocytes (graph: d-f, green bars), but these are significantly decreased by the end of first meiosis and not visible in metaphase-II-arrested oocytes. Minimum of three trials, except for e14.5 and primary follicles (two trials). Image panel: direct detection of GFP CETN2 doublets and singles after fixation and co-labeling with PCM antibody (y-tubulin or pericentrin; not shown) and DNA (not shown). Images are at similar exposure and size for comparative analysis. The drop in GFP signal background (F-H) reflects dilution of GFP signal as the oocyte volume increases during follicular growth. Bars = .4 μm.

Our analysis of the fate of GFP CETN2 centriole doublets, traced from e11.5 PGCs isolated from the genital ridge through mature metaphase-II-arrested oocytes is summarized in Figure 3 and Supplemental Figure S5. GFP CETN2 foci associated with y-tubulin or pericentrin PCM in PGCs, fetal oogonia, and early oocytes from P4 neonate ovaries showed double pairs of GFP CETN2 centrioles (Fig. 3a-c) closely apposed in the cytoplasm (Suppl Fig. S5a-c). Incompetent GVs (< 66μm diameter) from adult ovaries, as well as mature GV’s collected after hormonal simulation (> 75μm diameter), showed the average distance between double pairs of GFP CETN2 foci dramatically increased (Suppl Fig. S5d-e; <: p< 0.01), and one of the GFP CETN2 pairs became brighter (Fig. 3d). Interestingly, around 3-10% of incompetent or competent GV oocytes showed more than two doublet pairs (Fig. 3d-g, green bars). No evidence of centriole duplication in these quiescent oocytes was found, suggesting these aberrant oocytes were produced by fetal-stage mitotic errors or, perhaps, by early oocyte cyst breakdown inconsistencies. Often, oocytes with abnormal doublet pairs showed widely dispersed GFP CETN2 foci in the cytoplasm or cortex or along the GV surface, with distances > 12μm generally observed. Regardless, centriole dissolution appears to accelerate following meiotic resumption, with an increase in the number of single GFP CETN2 pairs at the spindle poles of metaphase-I and -II oocytes (Fig. 3, lower panel). Collectively, the data show that GFP CETN2 centrioles embedded within PCM are maintained as two doublets until follicular recruitment in post-adolescent females, when GV-arrested oocytes within growing follicles first separate into GFP CETN2 pairs and, later, single-pair dissolution occurs. Following meiotic resumption, single-centriole-pair dissolution accelerates. No evidence of centriole duplication was found during meiotic maturation. Of the metaphase-II-arrested oocytes with detectable GFP CETN2 foci at the spindle poles (∼ 38%; Suppl Fig. S2A), nearly all showed diminished single pairs at their spindle poles.

### IMMATURE, GROWING, AND MATURE OOCYTES CONTAIN GFP-CENTRIN FOCI IN γ-TUBULIN PCM

Fixed neonatal Day-4 post-birth female ovaries (P4), a stage at which centrioles have been identified by EM (Kloc et al, 2008), showed two tightly apposed doublets of GFP CETN2 foci within γ-tubulin PCM, adjacent to the GV (Fig. 4). Pre-antral and early antral follicles in adult ovaries maintained GFP CETN2 foci, with some displaying doublet pair separation, beginning with antral formation (Fig. 4C). Some cytoplasmic GFP CETN2 foci did not co-localize with y-tubulin or pericentrin PCM and were designated as assembled GFP aggregates.

**Figure 4.**
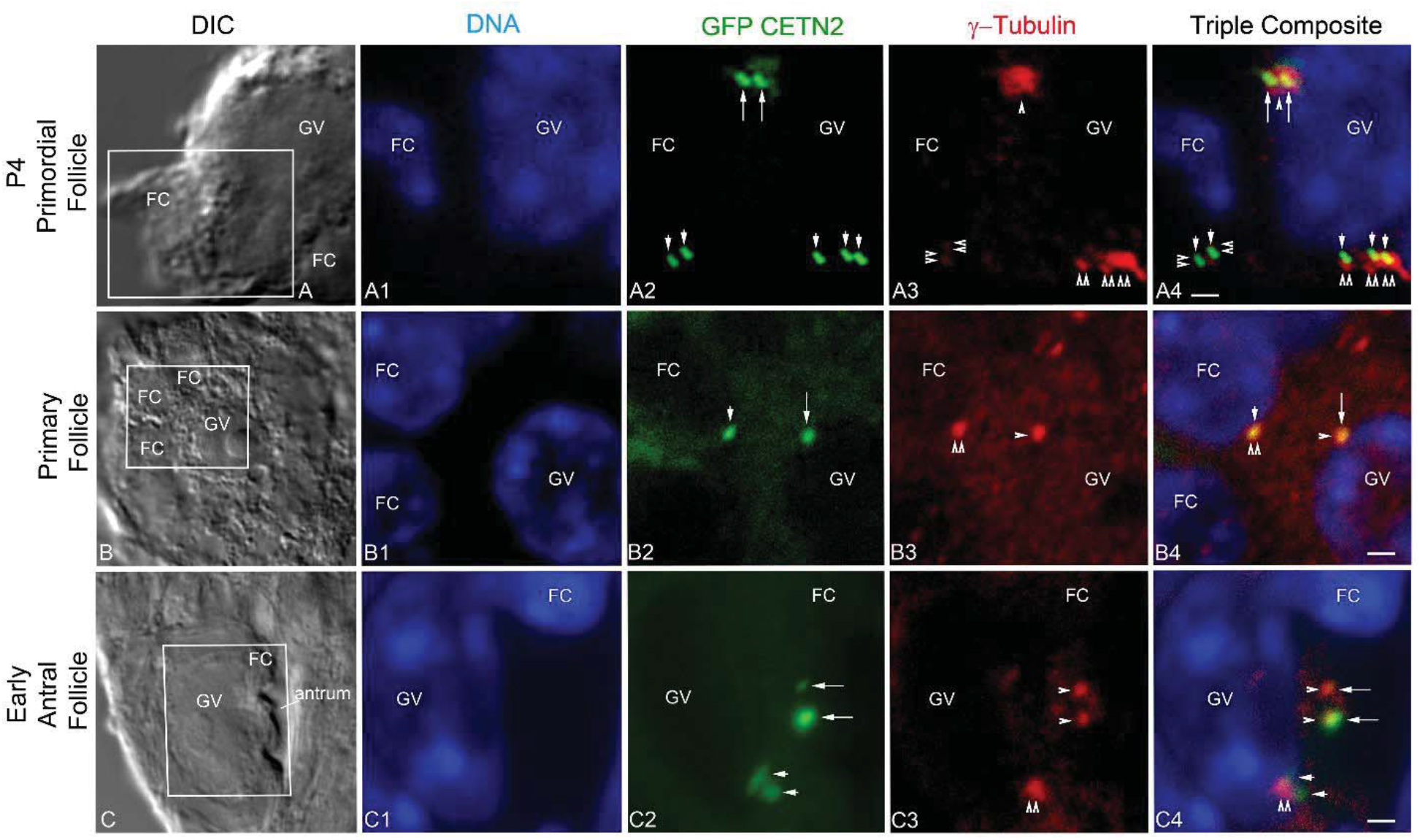
Immunohistochemical analysis of oocytes in GFP CETN2 ovaries from preadolescent P4 and adult mice. (**A1-A4**): P4 primary oocyte showing GFP CETN2 doublets (A2: green, long arrows) in association with the PCM protein pericentrin (A3: red, arrowhead) adjacent to the GV nucleus (A1: blue, DNA). Image also shows a somatic follicular cell GFP CETN2 doublet (A2: short arrows) embedded in pericentrin (A3: red, double arrowheads). A4: triple overlays of image panels A1-A3. (**B-B4**): A pre-antral follicle with a primary oocyte from an adult ovary showing a single GFP CETN2 foci (B2: green, long arrow) within y-tubulin (B3: red, arrowhead) at the GV nucleus (B1: blue, GV). A follicular cell (B1: blue, FC) with GFP CETN2 foci (B2: green, short arrow) in y-tubulin (B3: red, double arrowhead) is also visible. B4: triple overlay of image panels B1-B3. (**C-C4**): An early antral follicle from an adult ovary showing a GV oocyte with a pair of GFP CETN2 foci (C2: green, long arrows) within γ-tubulin (C3: red, arrowheads) at the nucleus (C1: blue, GV) and a surrounding follicular cell with GFP CETN2 centrioles (C2: green, short arrows) and γ-tubulin (C3: red, double arrowheads). C4: triple overlays of C1-C3 image panels. All images are 7-μm sections through ovaries taken from females expressing GFP CETN2 and counterstained with y-tubulin (red) and DNA (blue). Boxes in A, B, and C differential interference contrast (DIC) images are areas enhanced in the fluorescent image panels. Bars= 10μm.

Tracking GFP CETN2 doublet fate in ovarian sections is challenging and provides the rationale for isolating immature oocytes of assorted sizes during follicular growth (Suppl Fig. S6). Incompetent GV oocytes, which could not initiate meiotic maturation, showed GFP CETN2 foci embedded within γ-tubulin PCM in all oocytes, mostly without microtubule aster assembly (Fig. 5A). Only 70% of fully competent mature GV oocytes maintained GFP CETN2 foci with γ-tubulin (Suppl Fig. S2A). Although numerous GFP protein aggregates in growing GV oocytes were present, these supernumerary foci did not interact with γ-tubulin or pericentrin, nor assemble microtubules. The GFP CETN2 foci in association with γ-tubulin or pericentrin maintained consistent area sizes during oocyte growth, although their distribution within the oocyte changed from the GV nuclear surface in small follicle GVs to the cytoplasm during mid-follicular growth, before returning to the GV surface just before meiotic resumption (Suppl Fig. S2B-S2C).

**Figure 5.**
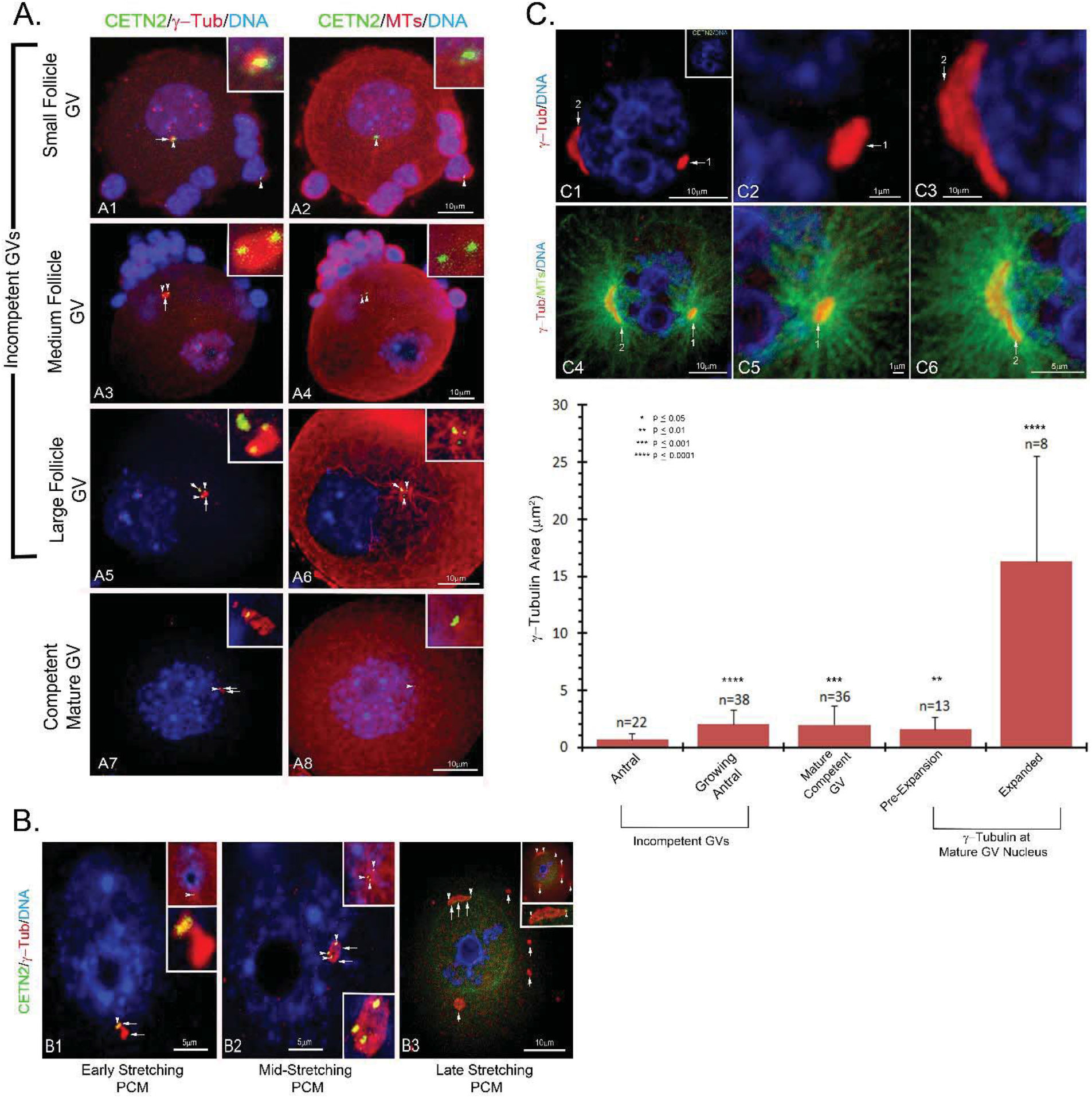
GFP CETN2 centriole characteristics during oocyte growth. (**A1-A8**): Selected confocal z-projections of follicle oocytes showing GFP CETN2 centrioles (green, arrowheads) and y-tubulin foci (red, long arrows) in pre-antral (A1, A2), growing incompetent (A3-A6), and mature competent GVs (A7, A8). Growing GV oocytes assemble cortical microtubule arrays rather than asters from the GFP CETN2-containing MTOCs (A2, A4, A8: red), with rare exceptions (A6: red). Cumulus cells have GFP CETN2 centrioles embedded in y-tubulin (A1, A2: solid arrowhead). Often, growing GV oocytes with abundant GFP CETN2 aggregates lacking y-tubulin assemble in the cytoplasm (A5, A6: short arrow). (**B1-B3**): Prior to meiosis resumption, the GV-residing MTOCs stretch on the nuclear surface (B1-B3: red, y-tubulin). GFP CETN2 foci (green, arrowheads) associate with the largest MTOC (B1-B3: red, long arrows), with the doublets splitting to opposite ends of the elongating MTOC (B3: green, arrowheads). Other MTOCs on the GV nuclear surface lacking GFP CETN2 foci have not expanded (B3: red, short arrows). (**C1-C6**): an oocyte taken from a GFP CETN2-expressing female lacking GFP CETN2 foci (C1-C3: green) in the MTOCs (C1-C3: red, y-tubulin, arrows). MTOC expansion on the GV nucleus still occurs as microtubule aster assembly increases (C4-C6: green, microtubules), indicating that MTOC expansion does not require embedded GFP CETN2 foci. (**Graph**): measured y-tubulin areas showing GFP CETN2-containing MTOCs significantly increase in size during oocyte growth to full GV maturation. Prior to nuclear envelope breakdown and meiosis resumption, GV MTOCs embedded with GFP CETN2 foci dramatically expand on the nuclear surface. N=number of y-tubulin focus areas measured. Key, upper left: *p values, determined by the two-tailed Student’s t-test (GraphPad Software). All images are GFP CETN2 oocytes triple-labeled for y-tubulin (A1, A3, A5, A7, B1-B3, C1-C6: red), microtubules (A2, A4, A6, A8, upper insets in B1-B3: cy5, color assigned red or C4-C6: color assigned green), and DNA (blue). A1, A3, A5, A7 insets: y-tubulin (red) and GFP CETN2. A2, A4, A6, A8 insets: microtubules (red) and GFP CETN2. B1-B3, lower insets: y-tubulin (red with GFP CETN2). C1 inset: GFP CETN2 (green) and DNA (blue). Bars, as marked.

Mature GVs, preparing to enter first meiosis, had decondensed, stretching PCM along the GV surface, vastly increasing the surface area (Fig. 5B and graph; Luksza et. al, 2013; Cliff and Schuh, 2015). Oocytes with two GFP CETN2 pairs typically separated to opposing ends of the enlarging, fragmenting MTOC (Fig. 5B3). Other nuclear and cytoplasmic PCM foci lacking GFP CETN2 foci did not immediately undergo expansion along the GV surface (Fig. 5B3, short arrows). However, PCM stretching along the GV surface does not require GFP CETN2 foci (Fig. 5C). Collectively, GFP CETN2 foci, present from early pre-antral follicle oocytes, are maintained through follicular growth to full maturity. These structures remain within the γ-tubulin or pericentrin PCM as it undergoes decondensation, stretching, and fragmentation along the GV surface in preparation for meiotic spindle assembly.

Since GFP CETN2 doublets appear to separate during GV growth, we investigated whether the expression of Cep135, a centrosomal scaffolding protein that anchors cNAP1 in somatic cell centrioles and whose disruption or overexpression causes premature centrosome separation (Kim et al, 2008), might be the underlying mechanism (Fig. 6). Adhering cumulus cells, as well as incompetent GVs isolated from small, medium, and large growing follicles, showed bright GFP CETN2 foci co-localized with pericentrin and Cep135 (Fig. 6A-F). As oocytes transitioned toward meiotic resumption, GV oocytes expanded and the MTOC fragmented along the GV surface as Cep135 detection was significantly reduced in the PCM and the GFP CETN2 doublet pairs separated (Fig. 6G-J). cNAP1, anchored by Cep135 at somatic cell centrioles, has been identified in mouse oocyte MTOCs (Sonn et al, 2010), and PLK1 has been implicated in release of cNAP1 to permit MTOC expansion in the mouse GV oocyte (Cliff and Schuh, 2015). Therefore, we utilized the PLK1 inhibitor BI 2536 to investigate whether the inhibitor impacts Cep135 localization at MTOCs and subsequent GFP CETN2 doublet separation. A 2-hr treatment of mature GV oocytes with BI 2536 inhibitor blocked Cep135 reduction from MTOCs at the GV surface, which still slightly expanded (Fig. 6K). Remarkably, despite Cep135 retention in the MTOC, the GFP CETN2 doublet pairs separated to opposite ends of the PCM (Fig. 6K-M). Even the combined treatment of BI 2536 with the microtubule inhibitor nocodazole, which blocks MTOC expansion and fragmentation, showed slightly elongated MTOCs on the GV surface, with retained Cep135 in the PCM and GFP CETN2 foci separated into four pairs (Fig. 6N-P). Collectively, these results suggest that GFP CETN2 doublet separation does not require the loss of Cep135 centrosomal core protein at the MTOC. Centriole doublet separation and dissolution may involve other unknown mechanisms occurring upstream of meiotic resumption.

**Figure 6.**
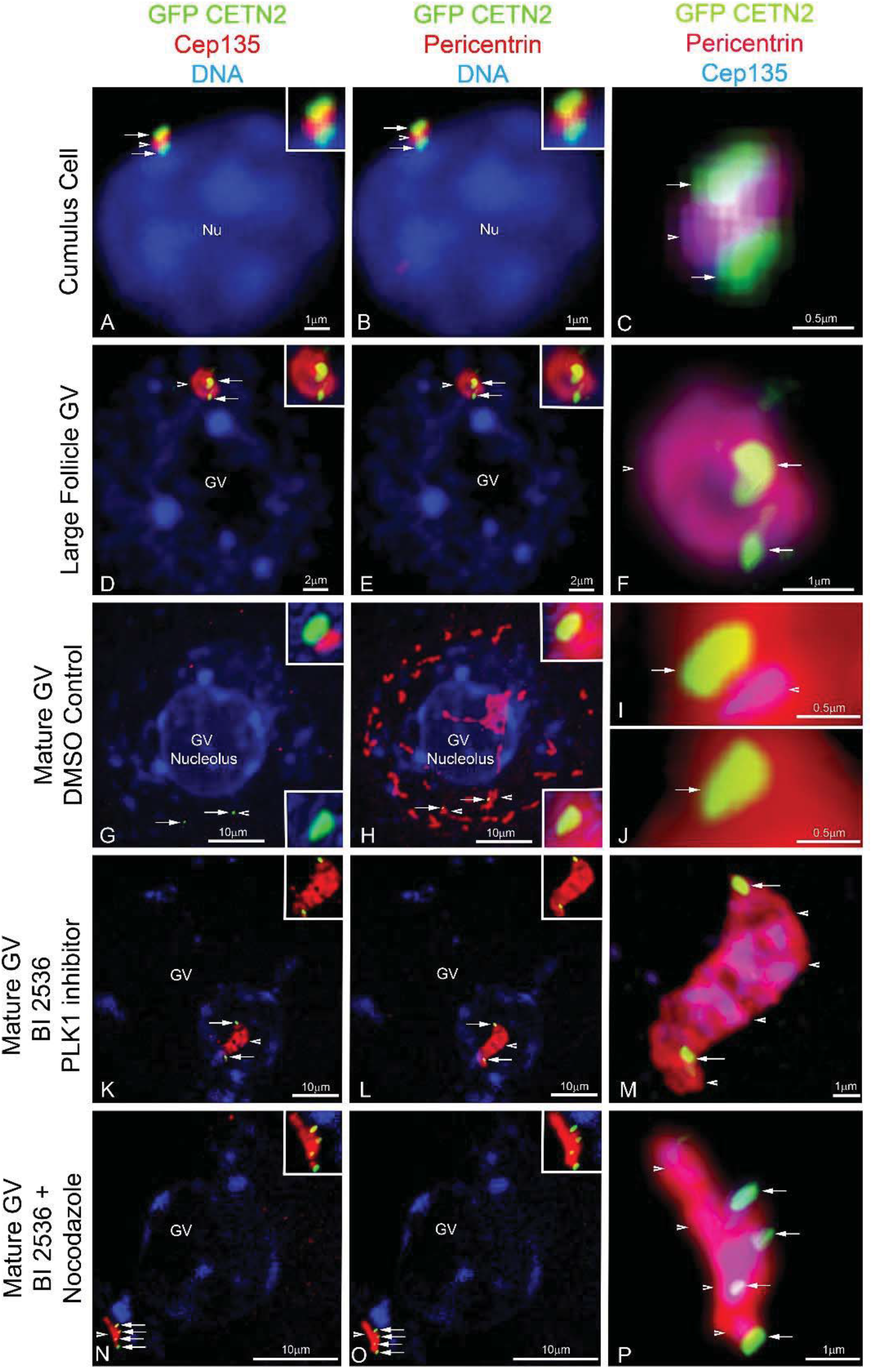
The loss of Cep135, a cNAP1-anchoring protein, may not be required for initial GFP CETN2 doublet separation. (**A-C**): a somatic cumulus cell (DNA, blue) with apposed GFP CETN2 centrioles (green, arrows) co-localized with Cep135 and pericentrin (red, arrowheads). C: overlay of GFP CETN2 (green, arrows), Cep135 (blue), and pericentrin (red, arrowhead), showing details of Cep135 and pericentrin co-localization between the centriole doublets. (**D-F**): an incompetent GV containing two CETN2 doublets (green, arrows) embedded in an MTOC with Cep135 and pericentrin (red, arrowheads) at the GV surface (blue). F: triple overlay of Cep135 (blue), pericentrin (red), and the GFP CETN2 doublets (green, arrows), showing oocytes maintain Cep135 within the MTOC during growth. (**G-J**): a mature oocyte 2 hrs post-culture with extensive MTOC stretching and fragmentation (H: red) on the GV surface (blue, DNA). Cep135 is significantly reduced in the expanding PCM (G: red), and the GFP CETN2 doublets separate (green, arrows) within pericentrin ribbons (H: red, arrowheads). (I-J): enhanced details of Cep135 (green, arrows), pericentrin (red), and reduced Cep135 (blue) at the separated doublets. (**K-M**): a mature GV oocyte incubated 2 hrs in BI 2536 PLK1 inhibitor, showing reduced MTOC stretching and fragmentation (red, arrowheads) with retained Cep135 (K: red) and pericentrin (L: red). Despite Cep135 retention, the GFP CETN2 doublets separate to opposing MTOC ends (green, arrows). M: magnified view of GFP CETN2 (green, arrows), Cep135 (blue, arrowheads), and pericentrin (red) organization. (**N-P**): a mature GV co-incubated 2 hrs simultaneously in BI 2536 and the microtubule inhibitor nocodazole (10μM). The elongated MTOC retains highly co-localized Cep135 and pericentrin (red, arrowheads), and the GFP CETN2 doublets separate with the PCM (green, arrows). P: detailed view of Cep135 (blue), pericentrin (red), and split GFP CETN2 doublets (green, arrows). All images are triple-labeled for pericentrin (red), Cep135 (color assigned red in A, D, G, K, N; blue in C, F, I, J, M, P), and DNA (blue). GFP CETN2 (green) is by direct fluorescent detection. Insets: details of GFP CETN2 foci with Cep 135 (A, D, G, K, N) or pericentrin (B, E, H, L, O). Bars, as marked.

### GFP CETN2 FOCI PRESENT A LOSE ASSOCIATION WITH THE PCM

To verify that the GFP CETN2 foci observed in fully grown mouse oocytes were not an artifact due to fixation, we also followed them in living oocytes. These investigations, which were performed in France on similar and identical mouse strains, obtained independently, provide confirmation of the reliability of both the research resources and the experimental strategies. For these experiments, oocytes expressing mCherry-Plk4 (either from transgenic mice or from cRNA injection; Marthiens et al, 2013), a PCM marker (Bury et al, 2017), together with GFP or Venus-tagged CETN2 (also from either transgenic mice or cRNA injection) were utilized. No differences in behavior or localization were observed between cRNA and transgenic expression of these constructs (Fig. 7, compare A and D). Live imaging also revealed the presence of GFP CETN2 doublets in follicular cells surrounding the oocyte coming from transgenic GFP CETN2 mice (Fig. 7A upper panels) as well as GFP CETN2 foci co-localizing with the cortical PCM in incompetent oocytes (Fig. 7A lower panel). The distance between the two doublets increased while the major PCM foci moved toward the nuclear envelope (Fig. 7B), arguing that anchoring of the major MTOC to the nuclear envelope favored the building of forces able to perturb its internal architecture, as observed by Luksza et al (2013). This distance increased even further in competent oocytes able to resume meiosis (Fig.7C). This is fully consistent with observations made in fixed oocytes (Suppl Fig. S5). Importantly, as meiosis I progressed, live imaging showed that the association between GFP CETN2 foci and the PCM, labeled with mCherry-Plk4, was weak (Fig. 7D and Suppl Movie S1) at all stages of meiosis I spindle assembly (Suppl Movies S2 and S3). Foci of GFP CETN2 could be extruded into the first polar body (Fig. 7D and Suppl Movie S1) as also observed in fixed oocytes (Fig. S2F). Collectively, live imaging demonstrated the capacity of GFP CETN2 foci to retain association and organize the PCM, although this co-localization was progressively lost as the oocytes resumed meiosis.

**Figure 7.**
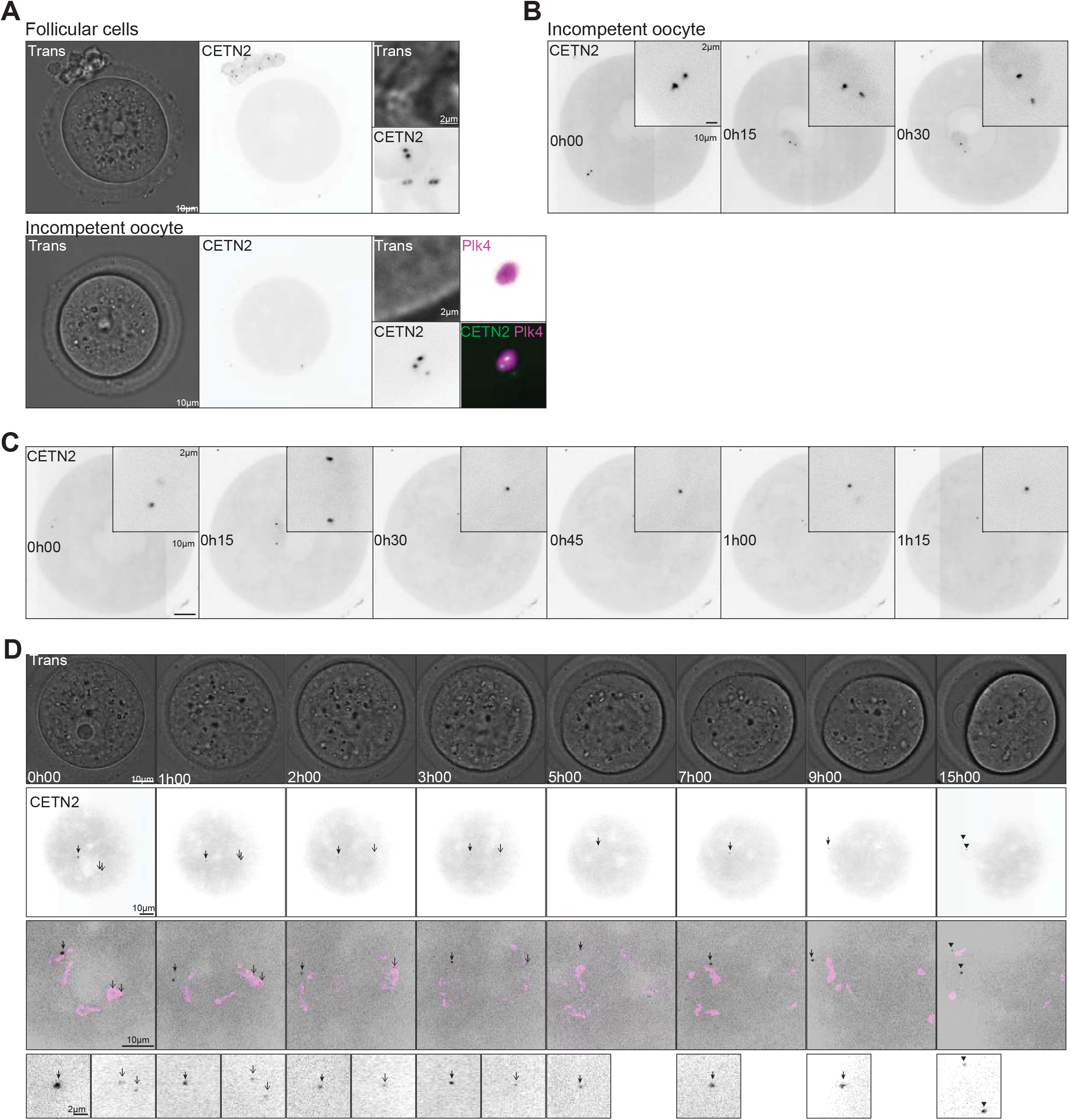
GFP CETN2 foci present a loose association with the PCM. (**A**): Transmitted light and fluorescent images of incompetent oocytes and follicular cells expressing GFP CETN2. GFP CETN2 signal (gray) is shown in follicular cells (upper panels) and incompetent oocyte (lower panels). Additional panels for the incompetent oocyte show the co-localization of GFP CETN2 (green) with mCherry-Plk4 (magenta). (**B**): GFP CETN2 foci (gray) moving from the cortex of the oocyte to the nuclear envelope in an incompetent oocyte. Insets show higher magnifications of the foci. (**C**): Oocyte followed at prophase I exit. GFP CETN2 appears gray. Insets show higher magnifications of the CETN2 foci. (**D**): Oocyte observed from prophase I exit to first polar body extrusion by transmitted light (upper panel) and fluorescence emission (lower panels). GFP CETN2 (gray) foci are highlighted with black arrows. Partial co-localization with mCherry-Plk4 (magenta) is shown. (**A-C**): GFP CETN2-expressing cells from transgenic mice, injected with mCherry-Plk4 cRNA (A). (**D**): mCherry-Plk4-expressing oocytes from transgenic mice, injected with Venus CETN2 cRNA. Time is expressed in hours and minutes.

### GFP CETN2 FOCI THAT FUNCTION IN MEIOTIC SPINDLE REASSEMBLY AFTER MICROTUBULE INHIBITION ARE PRESENT IN PCM

Since meiotic spindle assembly is initiated by multiple MTOCs, we explored whether GFP CETN2 foci within fragmented MTOCs would reassemble microtubules following disruption with the microtubule inhibitors nocodazole and paclitaxel (Fig. 8 and 9). GV-arrested oocytes incubated in the continuous presence of 10μM nocodazole for 3.5 hrs showed GFP CETN2 foci embedded in γ-tubulin PCM at the GV surface or in the cytoplasm (Fig. 8A-F). The GFP CETN2 MTOCs did not expand into ribbons, remaining as spheres despite the presence of nocodazole-resistant microtubules (Cliff and Schuh, 2015). A brief, 5-min rescue from nocodazole showed PCM increasing microtubule assembly prior to MTOC expansion (Fig. 8M-R). In prometaphase-I oocytes, 3.5-hr nocodazole exposure increased MTOC volumes near the condensed bivalents with embedded GFP CETN2 foci but no microtubule assembly (Fig. 8M-R). Upon a 5-min rescue from nocodazole, the MTOC with GFP CETN2 foci expanded and assembled a large, well-arrayed microtubule aster, suggesting that GFP CETN2-containing centrosomes can function as MTOCs that could potentially participate in spindle assembly (Fig. 8S-X).

**Figure 8.**
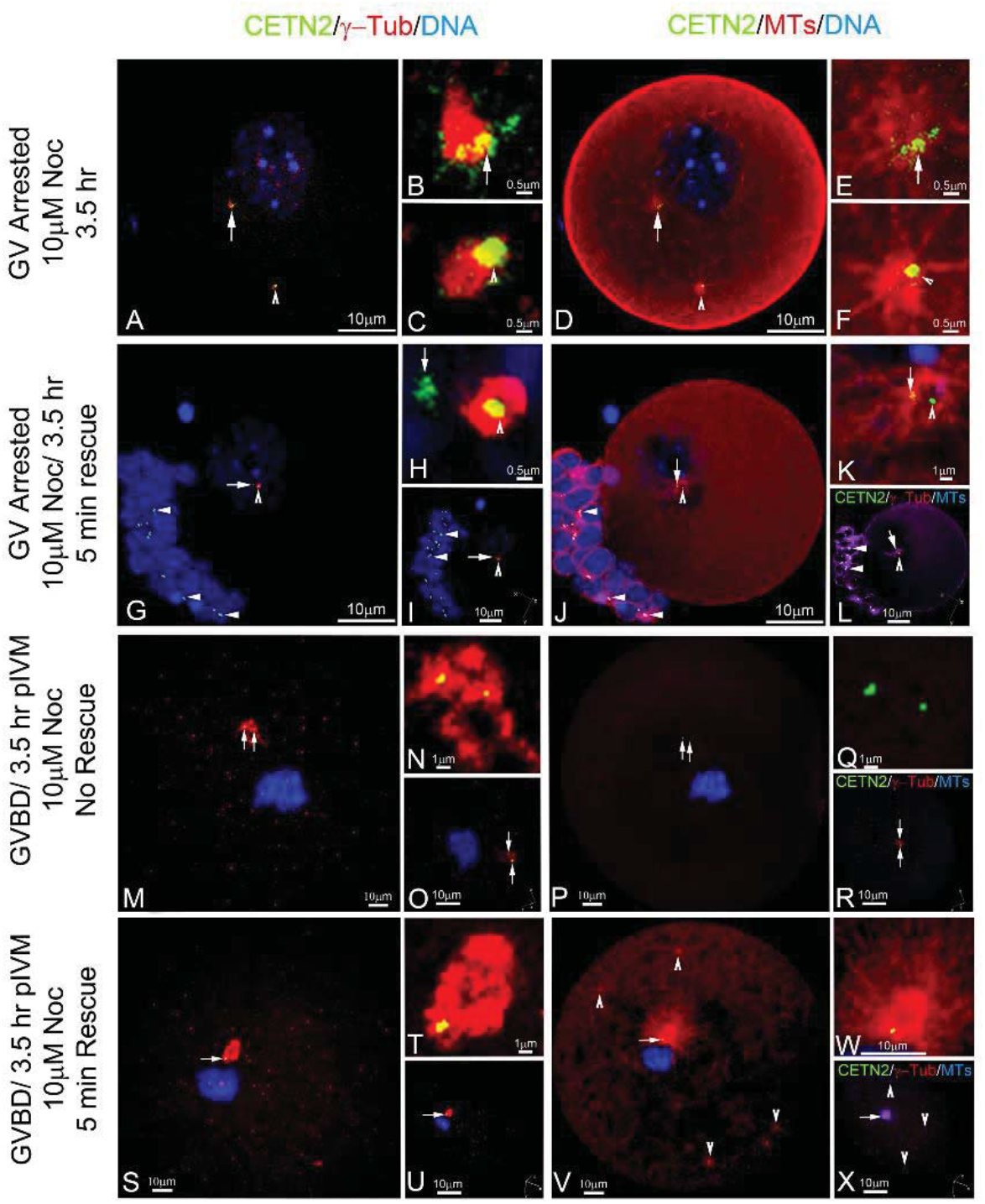
Recovery from nocodazole microtubule disassembly demonstrates functional GFP CETN2-containing centrosomes in GV and prometaphase-I oocytes. (**A-F**): a dbcAMP-arrested GV oocyte treated with 10μM nocodazole for 3.5 hrs shows one GFP CETN2 doublet at the GV (green, arrow) and a second in the cytoplasm (green, arrowhead). Both CETN2 foci associate with y-tubulin (A: red) and organize small nocodazole-resistant microtubule asters (D: red). B, C, E, F: details of GFP CETN2 (green) with γ-tubulin (B, C: red) or microtubules (E, F: red). (**G-L**): a GV-arrested oocyte treated as above but with a 5-min recovery from 10μM nocodazole inhibition. The GFP CETN2 foci (green, open arrowhead) are embedded within y-tubulin (G: red) at the GV (blue) and show microtubule aster assembly (J: red). G, J: red, arrows: a GFP aggregate lacking γ-tubulin lies within the microtubule aster. Cumulus cells (G, J: closed arrowheads) also have GFP CETN2 (green), y-tubulin (G: red), and cortical microtubules (J: red). H, K: details of GFP CETN2 foci (green, open arrowhead) and aggregate (green, arrow) in y-tubulin (H: red) or microtubules (K: red). (**M-R**): a GV oocyte treated with 10μM nocodazole for 3.5 hrs to produce a prometaphase-I oocyte with condensed bivalents (blue). GFP CETN2 doublets (green, arrows) reside in y-tubulin (red) around the bivalents (blue), but without assembled microtubules (P: red). N, O: details of GFP CETN2 in y-tubulin (N: red) or microtubules (Q: red). (**S-X**): an oocyte treated as in panel M but with a 5-min rescue from nocodazole clearly shows the GFP CETN2 doublets (green, arrow) in y-tubulin (S: red) rapidly assembling a microtubule aster (V: red; arrowheads, cytasters) at the bivalents (blue). T, W: details of GFP CETN2 foci in y-tubulin (T: red) or microtubules (W: red). I, L, O, R, U, X: oocyte image z-stacks in 3-D rotational view (axis, lower right) for GFP CETN2 (green), y-tubulin (red), and either DNA (I, O, U: blue) or microtubules (L, R, X: blue).

Paclitaxel enhances microtubule assembly. GV oocytes treated with lμM paclitaxel for 15 min did not undergo extensive MTOC fragmentation on the nuclear surface, and the GFP CETN2 foci remained at the core of the slightly decondensed PCM (Fig. 9A-F). Cortical microtubules were enhanced, especially around the GV, although microtubule aster assembly in the PCM containing the GFP CETN2 foci was not typically observed (Fig. 9D, red). After GVBD, paclitaxel initiated microtubule aster assembly around the GV nucleus from fragmented, dispersed MTOCs. The largest MTOC retained two embedded GFP CETN2 foci from which the best organized microtubule aster assembled next to the condensing bivalents (Fig. 9G-L). All other identified cytoplasmic and cortical microtubule asters assembled from MTOCs but without detectable GFP CETN2 foci (Fig. 9L). In later prometaphase-I, GFP CETN2 foci within many small fragments of γ-tubulin PCM were observed around the circular bivalents and extensive bundles of microtubules assembled outward toward the cortex in a large ring pattern (Fig. 9M-P). Taken together, paclitaxel-induced microtubule enhancement fragments MTOCs on the GV surface and prevents PCM assembly at opposing spindle poles. GFP-expressing CETN2 foci remain with the scattered MTOCs around the condensing bivalents, but these centrosomes neither nucleate microtubules nor assemble bipolar spindles.

**Figure 9.**
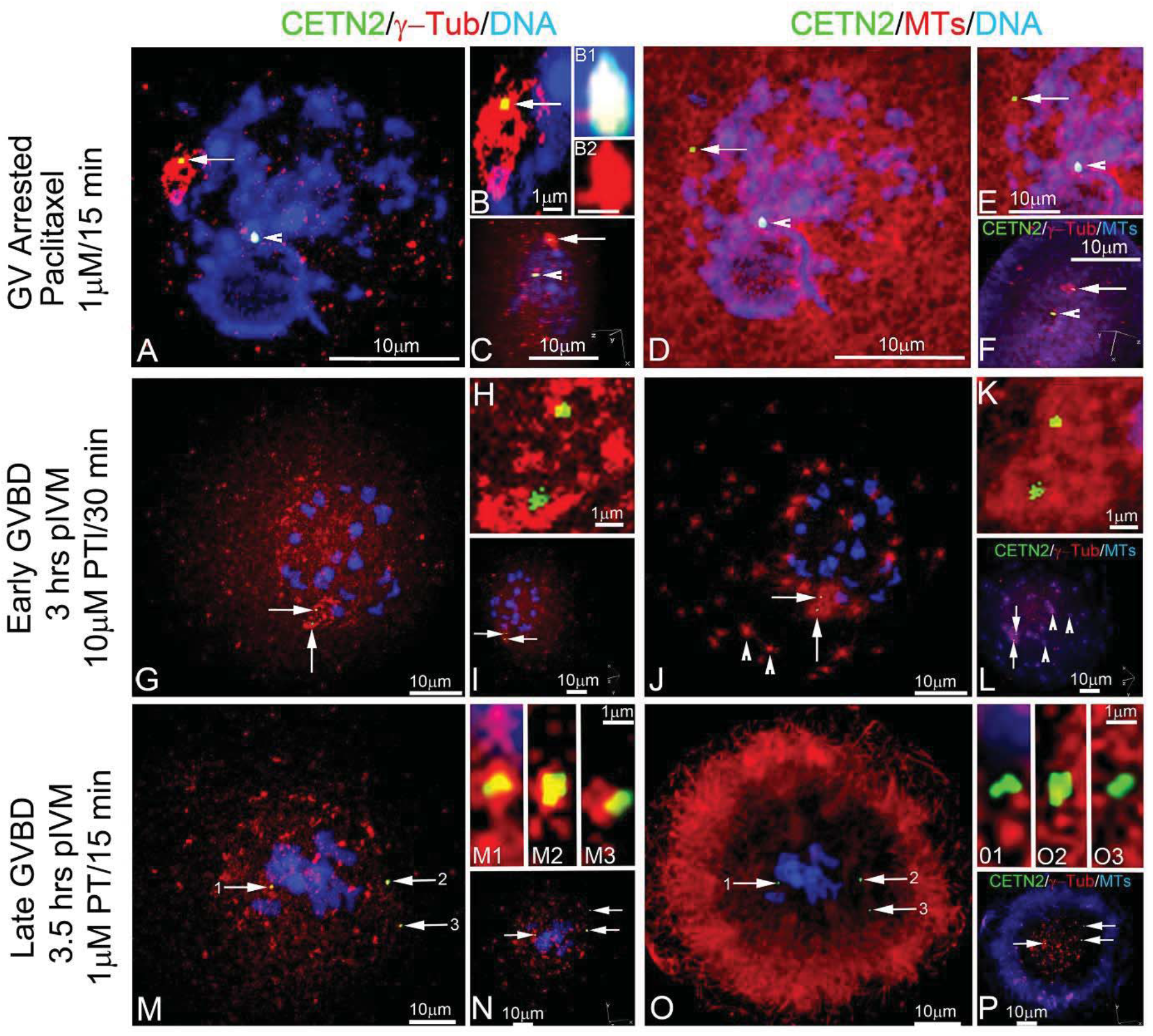
Paclitaxel (PT) enhances MTOC fragmentation around GFP CETN2-containing centrosomes after meiotic resumption. (**A-F**): A dbcAMP-arrested GV oocyte treated with 1μM PT for 15 min shows a GFP CETN2 doublet (green) embedded in y-tubulin PCM (red, arrow) on the GV nuclear surface (blue). A bright GFP aggregate at the GV surface is also observed (red, arrowhead). PT enhances cortical and cytoplasmic microtubules but not microtubule aster assembly at this stage (D: red). B, B1, B2: details of GFP CETN2 foci (B: red, arrow) or GFP aggregate (B1, green) with y-tubulin PCM (B: red). B2: nearby y-tubulin foci (red) adjacent to the GFP aggregate but not embedded in the PCM (see rotational view, C). (G-P): two GV oocytes cultured for 3.5 hrs to induce germinal vesicle breakdown before exposure to 1μM PT for 15 min. An early prometaphase-I oocyte (G-L) shows two GFP CETN2 doublets (green, arrows) in fragmenting y-tubulin (red) around condensing bivalents (blue). Enhanced microtubule aster assembly (J: red) is observed from the largest remaining y-tubulin PCM (G: red) having embedded GFP CETN2 centrioles (green, arrows). Abundant cortical cytasters (J: red, arrowheads) are also present after PT exposure. In later-stage prometaphase-I oocytes (M-P: blue), GFP CETN2 doublets separate (M: arrows) within the highly fragmented y-tubulin PCM (M: red). Microtubule assembly appears to radiate from the cytoplasm outward toward the cortex, not necessarily from the GFP CETN2 foci within y-tubulin PCM (O: red). H, K, M1-M3, and O1-O3: enhanced images of GFP CETN2 (green) in y-tubulin PCM (H, M1-M3: red) or microtubules (K, O1-O3: red). All images are GFP CETN2 oocytes counterstained with antibodies to y-tubulin (red), microtubules (D, E, J, K, O: color assigned red or F, L, P: color assigned blue), and DNA (blue). C, F, I, L, N, P: oocyte rotational 3-D volume views (axis, lower right), showing GFP CETN2 pairs (green), y-tubulin (red), and either DNA (C, I, N: blue) or microtubules (F, L, P: blue).

Analysis of changes to GFP CETN2 foci areas during exposure to nocodazole and paclitaxel is presented in Supplemental Figure S7. GV oocytes exposed continuously to nocodazole or paclitaxel for 3.5 hrs in the presences of dbcAMP to retard meiotic resumption showed significant decreases in overall GFP CETN2 foci areas, perhaps indicating GFP CETN2 foci instability in the face of altered microtubule dynamics (Suppl Fig. S7A, green bars). Similar observations were seen when dbcAMP was removed to permit GV oocytes to progress to prometaphase-I in the presence of nocodazole but not after paclitaxel exposure (Suppl Fig. S7A, blue bars). Regardless, GPF CETN2 foci areas decreased following resumption of meiotic maturation in all microtubule inhibitor treatments when analyzed with their corresponding GV controls (Suppl Fig. S7B). Collectively, disruption of microtubule assembly or disassembly reduces GFP CETN2 areas, even in arrested GV oocytes, suggesting that structural alterations have already occurred prior to oocyte full maturation that render the oocyte centrioles susceptible to changes in microtubule dynamics before meiotic resumption.

## Discussion

The transgenerational contribution of centrosomes and centrioles was thought to have been solved by Boveri (1887; Scheer, 2014) who wrote that: *“The ripe egg possesses all of the elements necessary for development save an active division-center (centriole). The sperm, on the other hand, possesses such a center but lacks the protoplasmic substratum in which to operate.”* Later, the pioneering ultrastructural imaging of centrioles by Fawcett and Phillips (1969), previously depicted as mere dots, recognized that centrioles and basal bodies were composed of 9-triplet microtubules. However, predictions that molecular genetic approaches would answer definitively inheritance questions, following on the discovery of DNA in mitochondria (Nass and Nass, 1963; Margulis, 1970), failed. Without a permanent tracer molecule, as DNA in mitochondria, it has not been possible to answer this question conclusively, until recently. Now, breakthroughs in understanding the nature and dynamism of the core protein components of both centrosomes and centrioles (Jana et al, 2014), and the realization that their assembly is tightly regulated and orchestrated provide the tools to address inheritance. With considerable simplification, centrioles assemble upon a SAS-6 cartwheel (Guichard et al, 2018), upon which the other components of the 9-triplet structures assemble and onto which centrosomal PCM later concentrates (Kollman et al, 2015). In consequence, the question as to whether centrosomes and centrioles are inherited, strictly speaking, may need to be examined more stringently. Findings demonstrating that they are elaborate, dynamic protein assemblies support the concept of “transmission,” as thoughtfully proposed by Ross and Normark (2015). Without the DNA of chloroplasts, mitochondria and other plastids, perhaps the term “inheritance” should not be applied casually to organelles, assembled at times *de novo*, which lack their own genomic material.

Centriole disassembly, studied here in oocytes during the last two meiotic divisions, remains challenging, though elimination mechanisms have been discovered in flies (Pimenta-Marques et al, 2016) and starfish (Borrego-Pinto et al, 2016). In mice, two centriole pairs persist from fetal stages (Lei and Spradling, 2016) and in oocytes within adolescent ovaries (Kloc et al, 2008). During maturation, the centriole pairs first separate within the surrounding PCM, with the disassembly of one of the two doublets prior to full oocyte growth. Centriole adhesion does not appear to depend on the Cep135 centriolar core protein that anchors cNAP1 at the centrioles. Centriole dissolution accelerates upon meiotic resumption, as the number and area significantly decrease following GVBD (Fig. 3; Suppl Fig. S2E). Perhaps these centriole-like remnants anchor the stretching PCM to the GV surface (Łuksza et al, 2013; Clift and Schuh, 2015). Later, the metaphase-I and -II spindles typically display at least a single pair near or at a spindle pole. Overall, this suggests that the maternal centriole is not lost until the completion of oocyte maturation, as in starfish oocytes (Borrego-Pinto et al, 2016; Verlhac, 2016). Centrioles were found in first or second mouse meiotic spindles neither by Szollosi, Calarco, and Donahue (1972) nor in our CLEM investigations here. Locating centrioles within an oocyte is akin to *looking for a needle in a haystack*, what with the enormous cytoplasmic volume and the lack of any fiduciary marks to help locate the centriole precisely. Here however, the CLEM approach defined the site of the GFP CETN2 pairs, and still, only osmiophilic PCM was observed (Suppl Fig. S1). This suggests that the canonical 9-triplet centriole structure disassembles prior to meiotic maturation, while the adjacent cumulus cells with intact centrioles provide ideal controls (Fig. 2E, inset and Suppl Fig. S1).

The GFP CETN2 doublets identified here likely fall into the category of centrioles described as “advertisements,” “remnants”, “passenger” or “zombie” structures (Mazia, 1987; Debec et al, 2010; Schatten and Simerly, 2015; Avidor-Reiss et al, 2015); i.e., vestigial, non-functional, and non-replicating centrioles. Perhaps centriole alterations mirror events seen in sperm distal centriole dissolution, where proximal centriole-like structures identified in the sperm of flies and humans lose their 3-dimensional 9-triplet microtubule structure, but retain PCM antigens and the ability to organize into functional MTOCs under the right conditions, such as after fertilization in the activated zygotic cytoplasm (Avidor-Reiss et al, 2015; Fishman et al, 2018). Indeed, the destruction of first one, and then the second, centriole in mouse sperm as these transit through the epididymis (Simerly et al, 2016) underscores the complex behavior of centrosomes and centrioles in gametes even after they depart from the gonad. While these GFP CETN2 doublets in maturing oocytes appear non-functional, their microtubule nucleating activity can be revived after recovery from nocodazole microtubule disassembly (Fig. 8). Upon this revival, robust microtubule arrays reassemble around the now reactivated GFP CETN2 doublets, which themselves, are surrounded by PCM foci.

The precise mechanism of CETN2 doublet separation is not currently known. Cep135, important in centriole biogenesis, is present in the MTOCs of incompetent oocytes and is lost when the PCM begins to expand and fragment prior to resumption of meiosis (Fig. 6). In somatic cells, Cep135 disruption leads to centrosome splitting and abnormal mitotic spindle phenotypes through its interaction with cNAP1, a centrosomal linker protein implicated in centriole-centriole cohesion (Mayor et al, 2000; Bahe et al, 2007; Inanç et al, 2013). cNAP1 is also in mouse GV MTOCs, although disruption of cNAP1 by microinjection of specific double-stranded RNAs did not affect first or second meiotic spindle assembly (Sonn et al, 2011). Recently, the initiation of MTOC stretching on the GV surface before nuclear envelope breakdown (Łuksza et al, 2013), critical for PCM fragmentation and first meiotic spindle assembly, was shown to be dependent on PLK1 phosphorylation of cNAP1, although the presence and functional role of cNAP1 in these MTOCs was unclear since they were not thought to have centrioles (Clift and Schuh, 2015). Our results suggest that maturing oocytes split the two CETN2 doublets early in follicular growth before attaining full maturation (Fig. 3), with Cep135 and cNAP1 continuing to be expressed until just before meiotic resumption. Thus, perhaps other proteins play a key role in mouse centriole-centriole cohesion upstream of Cep135 and cNAP1. Investigations on the precise molecules essential for centriole persistence and PCM functionality in the doublets and their fates during oocyte growth and maturation to metaphase-II arrest await future studies.

Centriole destruction may be prerequisite for terminal differentiation, and the retention of centrioles is associated with proliferation, regeneration, pluripotency, perhaps even totipotency. Skeletal muscle differentiation results in centriole loss (Tassin et al, 1985; Srsen et al, 2009) as during neuron differentiation (Li et al, 2017). Cardiomyocyte differentiation is particularly instructive, since centrioles are lost during terminal differentiation in murine hearts, which do not regenerate (Poss et al, 2002; Bettencourt-Dias et al, 2003). Remarkably, centrioles are retained in the regenerating hearts of fish and newts (Zebrowski et al, 2015). Here, centrioles are lost upon the onset of meiotic maturation, with immature oocytes retaining their double paired centrioles (Fig. 3, graph). Perhaps the last phase of oogenesis (i.e., oocyte maturation) is another example of terminal differentiation, including with the elimination of centrioles and centrosomes. Interestingly, while it appears necessary to undergo two mitotic cycles to generate centrioles, perhaps during both spermatogenesis and oogenesis, the last two meiotic cycles also are prerequisite for centriole destruction.

## Acknowledgements

We thank Bruce Campbell and Angela Palermo Lauff for expert editorial assistance and manuscript submission. We are grateful for the sponsorship of the National Institutes of Health and Pennsylvania Department of Health (#4100077065) for project funding. We also acknowledge the technical help of the core lab facility at Magee-Womens Research Institution for assistance with immunohistochemistry, as well as the important contributions of Dr. Alice Meunier (IBENS, Paris, France) for the gift of GFP-CENT2 mice and Dr. Renata Basto (Curie Institute, France) for the gift of the Venus-Centrin 2 construct and the mCherry-Plk4 mice. MHV acknowledges funding from the FRM (FRM Label-DEQ20150331758 to MHV) and the Inca (grant PLBIO 2016-270-TRAN) in France.

## Methods and Materials

### Mouse husbandry, handling, and Institutional oversight

Research in this study complied with the guidelines of the National Institute of Health’s Office of Laboratory Animal Welfare as described in the *Guide for the Care and Use of Laboratory Animals* regulations and by the DGRI in France (Direction Generale de la Recherche et de l’Innovation: Agrément OGM; DUO-1783). Our research protocols and ethics were approved by the University of Pittsburgh’s and Magee-Womens Research Institute’s Institutional Animal Care and Use Committees (IACUCs; protocol # 16027448). CB6-Tg (CAG-EGFP/CETN2)3-4Jgg/J mice (Stock number: 008234; Higginbotham et al, 2004) were derived from cryopreserved stocks at the Jackson Laboratory (Bar Harbor, ME) and hemizygous or wild-type genotypes acquired as juveniles. All mice were housed in a dedicated, Association for Assessment and Accreditation of Laboratory Animal Care-accredited mouse facility (light cycle: 12hr:12hr) or in the CIRB animal facility (French agreement number C75-05-12). Mice expressing enhanced GFP-labeled human centrin-2 transgene (designated GFP CETN2 CB6F1) were established by breeding 6-8-week-old hemizygous males or females with non-transgenic siblings or CB6F1/J inbred mice, and GFP CETN2 expression was confirmed by PCR or direct immunofluorescence, using tail tip tissue as previously described (Simerly et al, 2016). For this study, we investigated isolated germ cells from 14 gonads at e11.5, e14.5, and e18.5 as well as follicle and mature GVs from more than 36 females, producing more than 500 oocytes for analysis.

### Fetal stage sex determination by PCR

Genomic DNA was isolated for detection of the Y-chromosome in GFP CETN2-expressing e11.5, e14.5 and e18.5 mouse tail tip tissues and PCR performed using MyTaq Extract-PCR Kit (Bioline, Taunton, MA). A 331-bp segment of the gene was amplified using the Jarid primers (forward: 5’ CTG AAG CTT TTG GCT TTG AG; reverse: 3’ CCG CTG CCA AAT TCT TTG G; Life Technologies, Carlsbad, CA). B-actin served as a loading control, using the primers: 5’ GAT GAC GAT ATC GCT GCG CTG GTC G 3’ (forward), and 5’ GCC TGT GGT ACG ACC AGA GGC ATA CAG 3’ (reverse). *PCR* conditions were: 95°C for 3 min, followed by 35 cycles of 95°C for 15 seconds, 60°C for 15 seconds, and 72°C for 20 seconds. PCR product was analyzed in 2% agarose gel and visualized using ethidium bromide. Lanes expressing two bands were classified as males, with the remainder being female. Direct immunofluorescence was also employed to determine GFP CETN2 expression from non-gonadal cells at e11.5, e14.5, and e18.5 stages as previously described (Simerly et al, 2016).

### Fetal germ cell collection

Under IACUC-approved protocols, fetal gonads from GFP CETN2-expressing females were collected following mating with non-GFP males as described above. A copulation plug identified the morning after pairing was designated as 0.5-day post-coitus (dpc). Pregnant GFP CETN2 females were sacrificed at e11.5, e14.5, e15.5, and e18.5 dpc and fetal gonads harvested by the methods of DeMiguel and Donovan (2003). Briefly, implantation sites were isolated from uteri and embryos released after cutting decidua and applying pressure at the base with fine forceps. Placenta and amnion were mechanically removed from the embryos and a cut made below the fore limbs to remove heart, liver and intestines. The genital ridges, hindgut, and descending aorta were then removed intact before dissecting the gonads away from the descending aorta, gut, and mesonephros in sterile PBS. The isolated gonads from each embryo were placed in a 1.5-ml Eppendorf tube in 100μl PBS on ice until all dissections were completed. To release germ cells from the gonads, most of the PBS was replaced with 200μl of 0.05% trypsin:0.02% EDTA (trypsin, EDTA: ATCC, Manassas, VA) and incubated at 37°C for 8-10 min. Trypsin enzyme was inactivated with Trypsin Neutralization Buffer (ATCC) for 10 min at 37°C. After removal of neutralization buffer, the digested gonads were incubated in 200μl of warm modified human tubal fluid (HTF) medium without calcium or protein supplementation [mHTF-Hepes: 97.8mM NaCl/4.69mM KCl/0.20mM MgSO_4_▪7H_2_O/0.37mM KH_2_PO_4_/4.0mM NaHCO_3_/21.0mM HEPES/2.78mM glucose/21.4mM sodium lactate/100U/ml penicillin/100μg/ml streptomycin/5mg/liter phenol red; formulation modified from Irvine Scientific, Santa Ana, CA]. Gonads were mechanically pipetted 25-50 strokes with a P-200 sterile micropipette tip to release individual germ cells before fixation (see below).

### Follicular and mature germinal vesicle (GV)-stage oocyte collection

For follicle collection, GFP CETN2 CB6F1 ovaries were excised into warm (37°C) sterile EmbryoMax M2 culture medium (EMD Millipore, Billerica, MA) and minced mechanically, using sterile forceps. Follicles were segregated according to sizes (Suppl Fig. S6) and GV’s released using two sterile needles. Most adhering cumulus cells were removed by a mechanical pipet equipped with a sterile 75-μm tip (Stripper pipet; Origio Mid-Atlantic Diagnostics, Trumbull, CT). Diameters for each live oocyte were determined after capturing images using a Nikon digital sight camera (DS Fi1) with Elements software (Nikon USA, Melville, NY) for oocyte classification. All oocytes were maintained in KSOM until fixation. Mature oocytes were harvested from superstimulated ovaries induced by an intraperitoneal injection of 7.5 IU PMSG (Sigma-Aldrich, St. Louis, MO) for 48 hrs and mature oocytes kept inM2 culture medium supplemented with 100μg/ml dibutyryl adenosine 3’,5’-cycl¡c monophosphate (dbcAMP; Sigma-Aldrich) to prevent meiosis resumption. To initiate meiotic maturation, mature GVs were rinsed 3x in M2 culture medium without dbcAMP and placed in KSOM medium at 37°C in a 5% CO_2_ humidified incubator until fixation at the appropriate developmental stage.

### Cytoskeletal and PLK1 inhibitors

Nocodazole, paclitaxel and BI 2536 PLK1 inhibitors (Sigma Aldrich) were prepared as 10mM stocks in DMSO and stored in small aliquots at −80°C. All inhibitors were diluted to final concentrations in KSOM culture medium. Rescue experiments from cytoskeletal inhibitors were performed by rinsing treated oocytes 3x in M2 culture medium and returning to inhibitor-free KSOM for recovery at the times specified.

### Immunocytochemistry

Isolated primordial germ cells and oogonia from fetal gonads were attached to polylysine-coated (2 mg/ml; Sigma-Aldrich) 22mm^2^ coverslips and fixed in 2% paraformaldehyde (pFA; EM grade, 16% solution; Electron Microscopy Services, Hatfield, PA) in mHTF-Hepes without calcium or protein supplementation for 10 min at 37°C. After fixation, coverslips were rinsed extensively in PBS (no detergent) with 0.5% goat serum, blocked 30 min at room temperature in BlockAid solution (Thermo Fisher Scientific, Pittsburgh, PA), and primary antibodies applied simultaneously overnight at 4°C. For GFP CETN2 follicular and mature GVs, the zona pellucidae were first removed by 35-to 45-sec incubation in warm EmbryoMax acid Tyrode’s culture medium (Millipore) before the zona-denuded oocytes were attached to polylysine-coated coverslips and fixed in 2% pFA for 30 min at 37°C or 0.5% pFA (5 min), followed by postfixing for 5 min in absolute ethanol (−20°C). After fixation, coverslips were rinsed extensively in PBS with 0.25% Triton X-100 detergent (PBS-Tx), permeabilized in PBS + 2% Triton X-100 for 30 min at room temperature and then blocked 30 min in BlockAid solution before primary antibody application at 4°C overnight. GFP CETN2-expressing centrioles were detected by direct immunofluorescence. Primary antibodies to detect pericentriolar material, centriole antigens, germ cell markers, and microtubules included: mouse anti-SSEA-1 (Developmental Studies Hybridoma Bank, Iowa City, IA; 1:50), rabbit anti-Ak15 anti-γ-tubulin (Sigma-Aldrich; 1:500), mouse anti-pericentrin (clone 30; BD Transduction Laboratories, San Jose, CA), rabbit anti-Cep135 (Sigma Aldrich; 1:800), rabbit anti-Cep250/cNAP1 (Proteintech, Rosemount, IL), and rat anti-tubulin YOL 1/34 (EMD Millipore; 1:200), all diluted in PBS. Following overnight immunostaining, excess primary antibody was removed by several washes with PBS (for germ cells) or PBS-Tx (for oocytes) over 30 min, followed by application of appropriate fluorescently-tagged mouse, rat, or rabbit IgG or IgM secondary antibodies (1:500; Life Technologies) for 2 hr at room temperature in the dark. After final rinses in PBS or PBS-Tx, the DNA was labeled with Hoechst 33342 (10μg/ml; 10 min) before mounting coverslips in ProLong Diamond antifade (Life Technologies) and sealing with nail varnish.

### Immunohistochemistry of intact ovaries for primary oocyte detection

For immunohistochemistry, P4 neonatal and adult excised ovaries were fixed *in toto* in 4% pFA (Electron Microscopy Services) in mHTF-Hepes overnight at 37°C. Fixed ovaries were washed 3x in sterile PBS, air dried at −20°C for 2-4 hr, and then embedded in optimal cutting temperature compound (Tissue-Tek; VWR, Bridgeport, NJ) prior to storage at −80°C. For processing, 7-μm sections were cut on a cryostat at −20°C (CM1850 UV cryostat; Leica, Buffalo Grove, IL), floated onto clean glass slides (Diamond White Glass 25 × 75 × 1mm, (+) charged microscope slide; MidSci, Valley Park, MO), and further stored at −80°C until staining. To detect direct GFP expression, slides were placed at room temperature for 5 min, and a pap pen was used to demark a well around the sections. Slides were treated for 10 secs with Surgipath O-Fix (Thermo Fisher), washed briefly in distilled H_2_O, and immunostaining was performed with rabbit Ak15 anti-γ-tubulin (Sigma-Aldrich; 1:500) and rat anti-tubulin YOL 1/34 (EMD Millipore; 1:200) for 2hrs at room temperature. After primary antibody removal with PBS-Tx, appropriate secondary anti-rabbit and antirat secondary antibodies were applied for an additional 2 hr at room temperature. DNA was counterstained for 20 min with 10-μg/ml Hoechst 33342 DNA before sections were dipped in distilled H_2_O, blot dried, and antifade added prior to sealing with a 22 × 40mm coverslip and nail polish.

### Equipment, Imaging, analysis, and settings

Imaging of fixed slides was accomplished with a Nikon A1 four-laser line confocal microscope equipped with Elements A1 Plus compact GUI acquisition software (version 4.20; Nikon USA). Images were collected at 1024 × 1024 size at ¼ frames per second, using a pinhole size of 79.2 μm and a z-depth of 0.25 μm through the entire oocyte, with a differential interference contrast (DIC) Plan Fluor x100 (1.3 NA) objective. We collected 5- x 12-bit depth images (nd2 files), using the same laser photomultiplier tube settings for each channel across specimens (5% laser power, except UV, for DNA imaging, at 10.24%) to facilitate comparison between PGCs, oogonia, and immature oocytes from follicles of assorted sizes or for comparing control versus cytoskeletal inhibitor treatments. Fluorescent intensity ratios, mean intensities, area or volume measurements were performed on binarized images, using the threshold tool and region-of-interest statistical menu in the Elements software, and downloaded to Microsoft Excel for statistical analysis. For image panel presentation, we took the generated confocal nd2 files, subtracted a background image collected from outside of the oocyte, and then applied the deconvolution software module (Landweber; supplied by Nikon USA), using the point scan confocal command and same filter (clear) at eight iterations for all images. Final panels from deconvolved selected .tiff images were prepared in Adobe Photoshop (Adobe Systems, San Jose, CA).

### Correlative light and electron microscopy

CLEM was performed, as described previously (Kong and Loncarek, 2015), using GFP CETN2 mouse oocytes. Zona-denuded oocytes were attached on the glass surface of 35-mm Matek petri dishes coated with 2mg/ml polylysine and imaged at 37°C on an inverted microscope (Eclipse Ti; Nikon, Tokyo, Japan) equipped with a spinning-disk confocal head (CSUX Spinning Disk; Yokogawa Electric Corporation, Tokyo, Japan) to identify GFP CETN2 foci. After analysis by live imaging, Matek dishes were perfused with freshly prepared 2.5% glutaraldehyde, and 200-nm-thick Z-sections spanning the entire oocyte were recorded to register the position of GFP CETN2 doublets. Oocyte positions on the coverslips were then marked by diamond scribe, washed in PBS, and stained with 2% osmium tetroxide and 1% uranyl acetate. Samples were dehydrated and embedded in EMbed 812resin. The same oocytes identified by light microscopy were then serially sectioned. The 200-nm-thick serial sections were transferred onto copper slot grids, stained with uranyl acetate and lead citrate, and imaged using a transmission electron microscope (H-7650; Hitachi, Tokyo, Japan).

### Mouse strains and genotyping

Both CB6-Tg (CAG-EGFP/CETN2)3-4Jgg/J and ZP3 Cre+; mCherry-Plk4+ female mice were used. The [C57BL/6-Tg(Zp3-cre)93Knw/J] breeding pairs were obtained from Jackson Laboratories. mCherry-Plk4^flox/wt^ mice were generated by random insertion of a pCAG-loxCATloxmCherryPlk4SV40pA construct in the genome of C57BL/6N mice (Marthiens et al, 2013).

### Microinjection

Injection of *in* vitro-transcribed cRNAs into the cytoplasm of Prophase I-arrested oocytes was performed using an Eppendorf FemtoJet microinjector as described (Verlhac et al, 2000), and oocytes were kept for 1-3 hrs in milrinone to allow expression of fusion proteins. Oocytes were then released from their prophase I arrest by washing and transferring into milrinone-free M2 medium.

### Plasmid construction and in vitro transcription of cRNA

The mCherry-Plk4 construct used to produce the transgenic lines (Marthiens et al, 2013) and the Venus-Centrine 2 construct were subcloned into the pRN3 vector. cRNAs were synthesized using the T3 mMessage mMachine Kit (Ambion) and resuspended in RNase-free water (Verlhac et al, 2000).

### Confocal spinning disk imaging of living oocytes

Spinning disk movies were acquired using a Plan APO 40Å∼/1.25 N.A. objective on a Leica DMI6000B microscope enclosed in a thermostatic chamber set at 37°C (Life Imaging Services), and equipped with a CoolSnap HQ2/CCD-camera (Princeton Instruments) or EMCCD camera (Evolve) coupled to a Sutter filter wheel (Roper Scientific) and a Yokogawa CSU-X1-M1 confocal scanner. MetaMorph software (Universal Imaging) was used to collect the data.

### Statistics

Means ± standard deviations were determined by an online tool at EasyCalculations.com. We used Microsoft Excel to prepare graphs and box plots, which show median (horizontal lines), means (black squares), 25th and 75th percentiles (small boxes), and 5th and 95th percentiles (whiskers). Statistical significance was determined by Student’s *t* test (two-tailed), with actual p valves expressed, and was performed with GraphPad software (La Jolla, CA). Significance was determined at p < 0.05. Graphical analyses shown are indicative of average values ± standard deviation. For most experiments, more than three trials were performed, and data are representative of all trials.

### Data availability

Datasets generated and/or analyzed during this study are available from the corresponding author on reasonable request as set forth in the guidelines of this journal.

